# Stepwise Bayesian Phylogenetic Inference

**DOI:** 10.1101/2020.11.11.376459

**Authors:** Sebastian Höhna, Allison Y. Hsiang

**Affiliations:** GeoBio-Center, Ludwig-Maximilians-Universität München Richard-Wagner Straße 10, 80333 Munich, Germany; Department of Earth and Environmental Sciences, Paleontology & Geobiology, Ludwig-Maximilians-Universität München Richard-Wagner Straße 10, 80333 Munich, Germany

**Author notes:** To whom correspondence should be addressed: Sebastian Höhna, GeoBio-Center, Department of Earth and Environmental Science, Palaeontology & Geobiology, Ludwig-Maximilians-Universität München, Richard-Wagner-Straße 10, 80333 München, Germany, Phone: +49 (0)89 / 2180-6667.

**Keywords:** Bayesian inference, Joint posterior distribution, Parameter Uncertainty, Phylogenetics, Divergence time estimation, `RevBayes`

## Abstract

The ideal approach to Bayesian phylogenetic inference is to estimate all parameters of interest jointly in a single hierarchical model. However, this is often not feasible in practice due to the high computational cost that would be incurred. Instead, phylogenetic pipelines generally consist of chained analyses, whereby a single point estimate from a given analysis is used as input for the next analysis in the chain (*e*.*g*., a single multiple sequence alignment is used to estimate a gene tree). In this framework, uncertainty is not propagated from step to step in the chain, which can lead to inaccurate or spuriously certain results. Here, we formally develop and test the stepwise approach to Bayesian inference, which uses importance sampling to generate observations for the next step of an analysis pipeline from the posterior produced in the previous step. We show that this approach is identical to the joint approach given sufficient information in the data and in the importance sample. This is demonstrated using both a toy example and an analysis pipeline for inferring divergence times using a relaxed clock model. The stepwise approach presented here not only accounts for uncertainty between analysis steps, but also allows for greater flexibility in program choice (and hence model availability) and can be more computationally efficient than the traditional joint approach when multiple models are being tested.

## Introduction

Bioinformatics pipelines often contain multiple statistical analyses. Each statistical analysis is chained after the previous analysis by using the output from the first analysis as the input of the next analysis. However, in Bayesian inference, the perfect analysis would consist of a single analysis with a hierarchical model where each step of the analysis pipeline corresponds to one or more layers of the hierarchical model (Huelsenbeck et al. 2000b; Huelsenbeck and Rannala 2003). Performing such a multi-layered analysis is almost always impossible because of (1) the lack of software which implements such hierarchical models, and (2) the computational intractability of estimating all parameters simultaneously.

In phylogenetics, a chained analysis pipeline could consist of the following steps: (1) estimating a multiple sequence alignment for each locus; (2) estimating a gene tree from each locus; (3) estimating a species tree from the collection of gene trees; (4) estimating divergence times of the species tree; and finally (5) estimating diversification rates from the divergence times. Although it is possible in theory, there have yet to be any statistical analyses which perform all of these steps jointly, where all of these parameters are estimated in a single analysis. Instead, the usual bioinformatics approach is to perform each analysis step-by-step, using either Maximum Likelihood estimation or Bayesian inference.

Naturally, we can never be completely certain about our statistical estimates because there is always some associated uncertainty. For example, we will never know exactly the age of the split between humans and chimps; rather, we can only continue to narrow confidence or credible intervals with improved analyses. Unfortunately, this uncertainty is not propagated in a chained analysis pipeline because only a single estimate is used in the downstream analyses (though sometimes a small number of alternatives are tested). To address this, we introduce here a formal theory of *stepwise Bayesian inference*. Stepwise Bayesian inference is, in theory and in the limit, equivalent to *joint Bayesian inference*. While some variant of stepwise Bayesian inference has been used before in phylogenetic applications (*e*.*g*., Pagel and Lutzoni 2002; Pagel et al. 2004; Nylander et al. 2008), the equivalence between joint and stepwise Bayesian inference has not been established and explicitly tested before.

We introduce our stepwise Bayesian inference approach first using a toy example to verify the theory and demonstrate potential problems. Next, we demonstrate the utility of stepwise Bayesian inference for divergence time estimation using relaxed clock models. We show that divergence times obtained from joint and stepwise Bayesian inference are identical if sufficient information in the data exists and enough samples approximating the posterior distribution are available.

## Methods

### Stepwise Bayesian inference

The goal of any Bayesian statistical analysis is to estimate the posterior distribution of the parameter(s) of interest. Let us denote the parameter of interest as *θ* — for example, the phylogenetic tree with divergence times — and the data as *𝒟*. Bayes’ theorem gives us the posterior distribution of our parameters of interest **P**(*θ*| *𝒟*) as

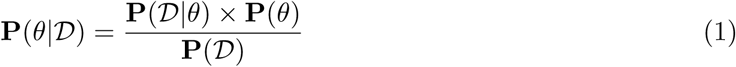

where **P**(*𝒟*|*θ*) is the likelihood function, **P**(*θ*) is the prior distribution of our parameters, and **P**(*𝒟*) is the marginal probability of the data (often simply called a normalizing constant).

Now let us assume that we have a slightly more complex model with additional parameters *ξ*, for example, the substitution model parameters (the exchangeability rates and the stationary frequencies) and the branch-specific clock rates. The posterior distribution of *θ* is then

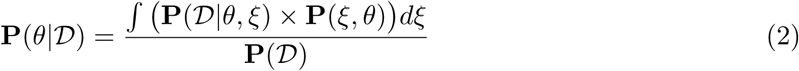

where we now need to integrate over all nuisance parameters *ξ*. In most cases, this integration cannot be done analytically and instead the model is augmented by these parameters. Standard numerical algorithms such as Markov chain Monte Carlo (MCMC) sampling (Metropolis et al. 1953; Hastings 1970) can be applied to integrate over all values of *ξ*. In this standard way we *jointly* estimate *θ* and *ξ*.

If our model has a hierarchical structure where **P**(*ξ, θ*) = **P**(*ξ*, |*θ*)×**P**(*θ*) and **P**(*𝒟* |*θ, ξ*) = **P**(*𝒟* |*ξ*), that is, *θ* are the parameters of the prior distribution on *ξ* and the probability of our data depends only on *ξ* but not *θ*, then we can rewrite Bayes’ theorem further as

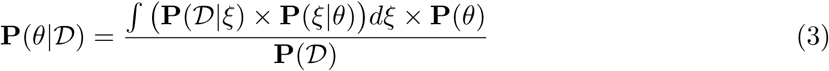

which we rearrange to

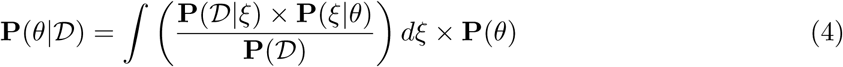

Next, we expand Equation 4 by multiplying both the numerator and the denominator with **P**(*ξ*), which is a prior distribution on *ξ* that is independent of *θ*. We thus obtain

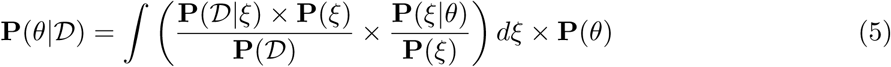

and realize that 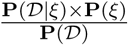 is the posterior distribution of *ξ* given the data. Thus, we further rewrite

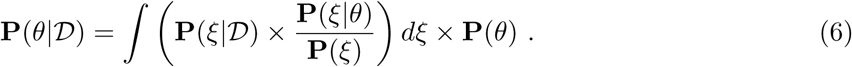

While this simplifies Equation 3, we are still left with the computationally difficult integral over the posterior distribution of *ξ*.

Our approach to avoid computing this integral adopts the standard sampling theory of distributions. As an analogy, assume that you want to compute the mean of a continuous distribution *f* (*x*), where the mean is defined in its common form as E[*𝒳*] = *∫x* × *f* (*x*)*dx*. Furthermore, assume that you cannot compute the integral over the distribution *f* (*x*) analytically. Nevertheless, what you can do, and most likely have done many times to compute the mean of a distribution, is to take a sample *𝒳* ∼ *f* (*x*) and compute the average of the sample as 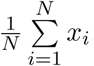. If you have sufficiently many samples, then the average of the sampled values will be a good approximation of the analytical mean without the need to compute the integral.

Hence, we adopt this *importance sampling* approach to avoid computing the integral over *ξ* in Equation (6). We generate *N* draws of *ξ* from the posterior distribution *P* (*ξ*| *𝒟*), which denotes the first step in our stepwise Bayesian inference approach. This allows us to replace the integral by a simple sum over the sampled values of *ξ*:

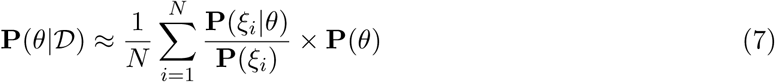

Equation (7) denotes the second step in our stepwise Bayesian inference approach.

Note that Equation (7) does not contain any terms using the data *D*. Therefore, computation of Equation (7) is generally faster than computation of Equation 4 or computing the posterior density *P* (*ξ*| *𝒟*), which involves the computationally expensive calculation of the likelihood **P**(*𝒟*|*θ*).

The fundamental assumption of our approximation of the integral over *ξ* is that a sample of *N* values is a sufficiently close approximation. If the posterior distribution **P**(*ξ*| *𝒟*), which we use as our importance distribution, is wide (*i*.*e*., has a large variance), then we will need more samples to approximate the integral well. Conversely, if this posterior distribution **P**(*ξ*| *𝒟*) is highly concentrated because there is a large amount of information in the data, then fewer samples will suffice. It is crucial to stress the importance of the number of samples in the first step because traditional bioinformatics pipelines use only a single value estimated from one step (*e*.*g*., the maximum likelihood estimate) as the input for the next step. We will test and demonstrate the effect of the number of samples on the accuracy of parameter estimation in the following sections.

### Multiple stepwise Bayesian inference

The models we present here have only two layers, *i*.*e*., with hierarchical structure **P**(*ξ, θ*) = **P**(*ξ*, |*θ*) × **P**(*θ*) and **P**(*D*|*θ, ξ*) = **P**(*𝒟*|*ξ*). However, the stepwise inference approach can be easily extended to multiple layers, as shown here in a three-layer model with additional parameters *ζ* (for example, diversification rate parameters):

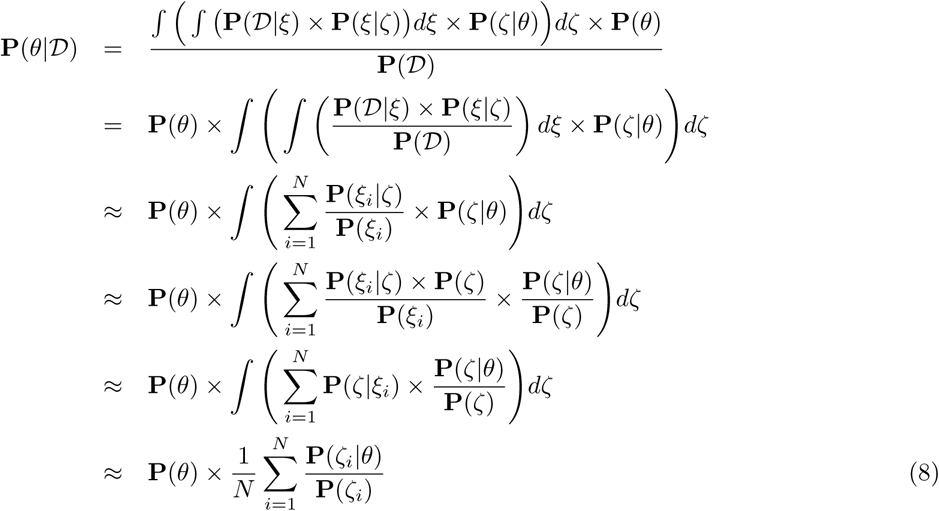

Equation 8 shows that it is possible and valid to divide each layer of the hierarchical model into a single step of stepwise Bayesian analysis. For each layer, we estimate the posterior distribution of the parameters and use this posterior distribution as the importance distribution in the next analysis. In this way, uncertainty in our parameter estimates is propagated from the first step through the last step.

### Comparison with previous stepwise Bayesian approaches

Previous work using a stepwise approach in a Bayesian phylogenetic framework differs from our approach in key ways: previous approaches apply only to combined data analyses instead of complex hierarchical models (Figure 1). Pagel et al. describe a similar 2-step method applied to comparative phylogenetics while accounting for uncertainty in the underlying phylogenetic tree (Pagel and Lutzoni 2002; Pagel et al. 2004). The major differences between the Pagel et al. stepwise approach and our stepwise approach can be summarized by graphical models of their overall respective structures, as shown in Figure 1. Specifically, our approach assumes the phylogenetic model has a hierarchical structure, that is, the probability of a given layer in the model is dependent only on the distribution of the parameter(s) in the layer immediately preceding it (Figure 1a). Given this hierarchical structure, we can treat samples from any given layer as the data for the following layer. For example, the phylogenetic model we present has two layers: (1) the layer where we infer the posterior distribution of the unrooted branch length trees (corresponding to *step 1* ; green box in Figure 1a); and (2) the layer where we infer the rooted tree with divergence times using an importance sample from the posterior inferred in *step 1* (corresponding to *step 2* ; coral box in Figure 1a). In this way, we sum over all of the importance samples in *step 2* and compute the average likelihood given the samples as data.

**Figure 1:**
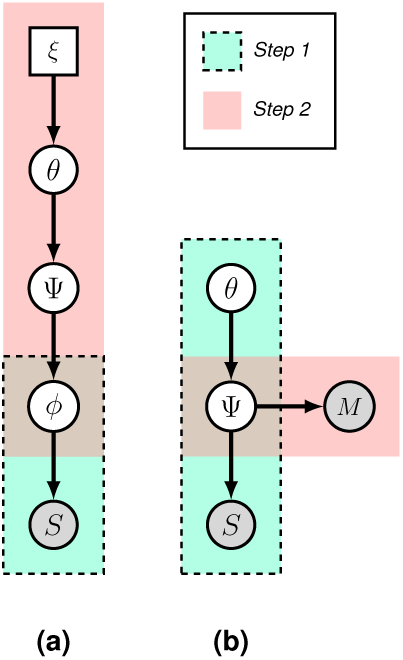
Graphical models showing difference between our stepwise approach (a) and the Pagel et al. stepwise approach (b). The components of the model involved in *step 1* for each approach are boxed in green with a dashed outline, and the components involved in *step 2* are boxed in coral. Our approach uses a hierarchical structure where an importance sample of the estimates from a preceding layer is used as data for the immediately succeeding layer. In contrast, the Pagel et al. approach assumes conditional independence between datasets and uses the sample of trees from *step 1* as a prior for the second step. Here, *S* refers to DNA sequence data, *M* to morphological data, Ψ to rooted ultrametric trees, *ϕ* to unrooted branch length trees, *θ* to the tree parameters, and *ξ* to the evolutionary model parameters.

In contrast, the Pagel et al. approach assumes a combined data analyses of multiple datasets (Figure Fig. 1b). That is, the sequence data (node S) and the morphological trait data (node M) are used consecutively instead of jointly. The first step of the Pagel et al. approach is to infer trees using the sequence data (green box in Figure Fig. 1b). In the second step (coral box in Figure 1b), a sample of the trees inferred during *step 1* is used as a prior for *step 2*, and a special MCMC move over the trees is used to integrate over these samples and compute the likelihood of the morphological trait data. Note that one could also sum over all tree sampled in *step 1* instead of using s specialized MCMC proposal (see our derivation below).

The Pagel et al. approach thus sums over data, whereas our approach sums over parameters. Our assumption is that the importance distribution is a good approximation of the posterior distribution of the parameter we are integrating over; this assumption is corroborated by our toy model results (see Figure 2b). Pagel et al. ‘s assumption is that the posterior distribution of the first step (*e*.*g*., using only genetic sequence data) is a good approximation of the posterior of the full joint analysis (which would include the morphological data).

**Figure 2:**
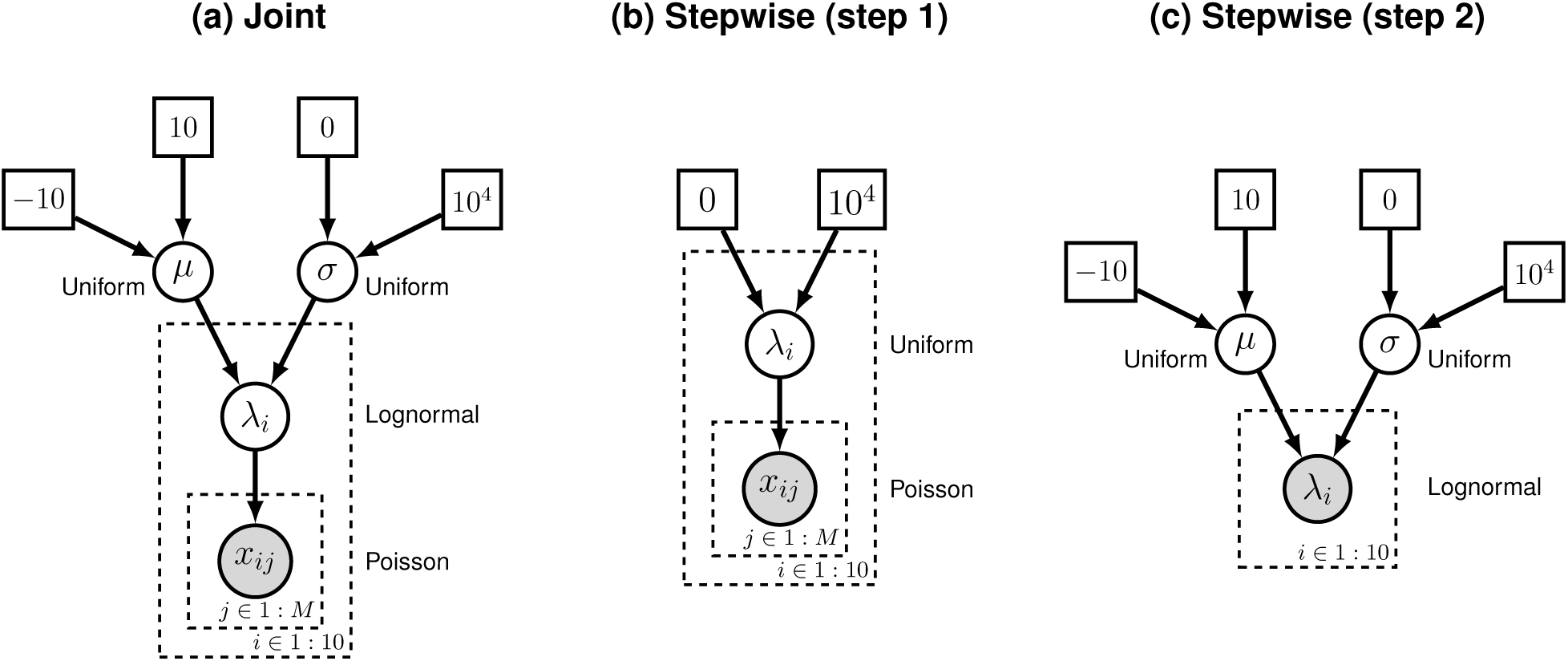
Graphical model of the (a) joint and (b,c) stepwise Bayesian inferences for the simple example. *M* observations *x*_*ij*_ were drawn from each Poisson distribution with parameter *λ*_*i*_. We arbitrarily picked 10 such Poisson distributions. Each *λ*_*i*_ was drawn from a lognormal distribution with parameters *μ* and *σ*. Both *μ* and *σ* had uniform hyperprior distributions. (a) The joint Bayesian inference uses a hierarchical model and estimates all *λ*_*i*_, *μ* and *σ* jointly. (b) The stepwise Bayesian inference first estimates the importance distribution on each *λ*_*i*_ (*step 1*); then, (c) the posterior distributions on *μ* and *σ* are estimated using samples from the importance distribution on the *λ*_*i*_s as observations (*step 2*).

### Derivation of the equivalence between stepwise Bayesian approach and joint Bayesian approach for multiple data sources

Let us assume we are interested in the rate parameter *κ* of our comparative model. Furthermore, we want to take the uncertainty in the phylogenetic tree into account, that is, we want to integrate out the phylogeny. We have two sources of data: the molecular sequence alignment *S* and the morphological character data *M*. The posterior probability of the rates *κ* are defined as

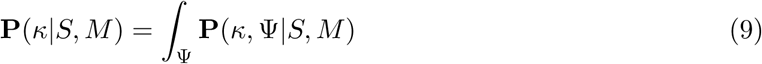

which we can expand and write in it’s full form as

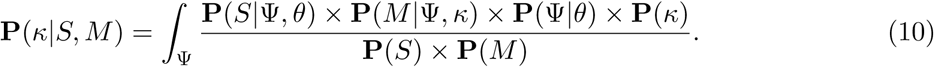

Equation 10 represent the full joint posterior probability of the rate parameter *κ*. Now we wish to split the joint analysis into a stepwise analysis. Therefore, we separate the equation into two parts. The first part represents the model of the molecular data and the second part represent the model of the morphological data,

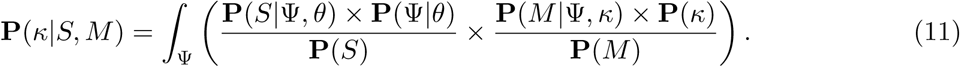

We then realize that the first part is equivalent to the posterior distribution of the phylogeny Ψ given the molecular sequence data *S*, and rewrite the equation as

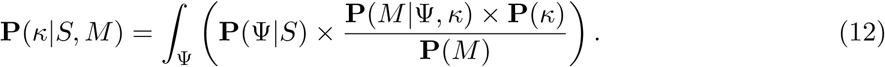

Thus, we perform *step 1* of the stepwise Bayesian analysis to estimate the posterior distribution of phylogenies given the molecular sequence data, **P**(Ψ|*S*). That is, we generate a random sample of *N* phylogenies from the posterior **P**(Ψ|*S*). Finally, we want to replace the integral over all phylogenies (tree topologies and branch lengths) by this importance sample frpm **P**(Ψ|*S*). Then, we can compute the posterior probability of the rates of morphological evolution *κ*

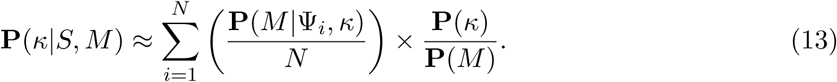

which represent *step 2* of the stepwise Bayesian inference.

We can conclude a few important facts from Equation 13. First, it is mathematically valid to use the samples of the phylogeny taken posterior distribution given the molecular sequence data (*step 1*) and compute the posterior distribution of the comparative model parameters using the average comparative model likelihood over all sampled trees. Second, the derivation follows exactly the same rational and motivation as our stepwise Bayesian approach for hierarchical models. Thus, we expect that the same properties apply, that is, many samples from the importance distribution are needed if (a) the information in the data in *step 1* is low, or (b) if the importance distribution and the joint posterior distribution differ significantly. Third, these derivation are independent of the actual probability distributions and models. Therefore, such a stepwise Bayesian analysis could be applied to any sequence of independent datasets.

### Branch-Rate-Tree Prior

The formulation of the branch-rate-tree prior requires a transformation between the two branches that subtend the root in the timetree to the corresponding single branch in the unrooted branch-length tree. Here we describe the transformation and the calculation of the Jacobian factor necessary to correctly calculate the likelihood under the branch-rate-tree prior.

Let us denote the branch length left of the root with *x* and the branch length right of the root with *y*. Similarly, let us denote the branch length left of the root in units of time as *t*_*x*_ and right of the root *t*_*y*_. Furthermore, assume any probability density on branch rates 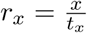 and 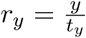, for example, a lognormal distribution with parameters {*μ*_*x*_, *σ*_*x*_} and {*μ*_*y*_, *σ*_*y*_} respectively:

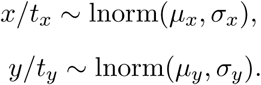

Now let us consider the transformation *u*, which is the sum of the left and right branch lengths, and the root fraction *v* as

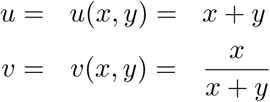

with its reverse transformations

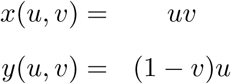

Next, we need to compute the derivates

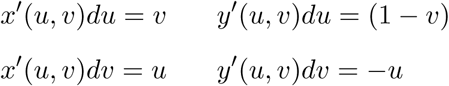

Therefore, we get the matrix *M* as

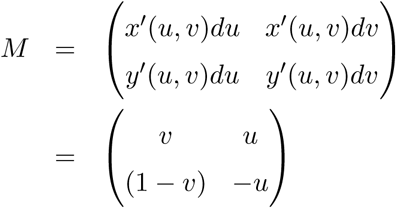

Finally, we can compute the Jacobian as

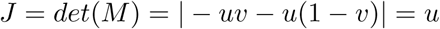

Hence, the absolute value of the Jacobian is the sum of the left and right root branch, or in other words, the root branch. Therefore,

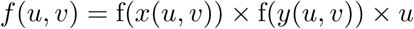

in the general case, and

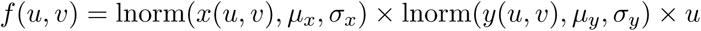

if we assume a lognormal branch rate prior distribution as an example.

### Implementation

We implemented the stepwise Bayesian approach in the software `RevBayes` (Höhna et al. 2016). The specific features presented here, especially the new *branch-rate-tree* distribution, are available from version 1.1.1 or later. The propagation of uncertainty is achieved by the `EmpiricalSample` distribution, which takes a sample of data instead of a single data element. This `EmpiricalSample` distribution can receive any distribution as its base distribution and thus provides a general way for any type of hierarchical model, not only the relaxed clock analysis as exemplarily used here. The source code is available from https://github.com/revbayes/revbayes. A tutorial can be found at https://revbayes.github.io/tutorials/step bayes.

The stepwise approach has also been previously used for phylogeographic applications, *e*.*g*., sequential Bayesian phylodynamic inference (Faria et al. 2014). This approach (implemented in the `BEAST` software suite using the `empiricalTreeDistributionModel` option) is equivalent to the approach of Pagel et al. only applied to biogeographic characters rather than phenotypic traits, and is also limited to distributions of phylogenetic trees. In contrast, the generalized theory presented here, combined with the flexibility of the `RevBayes` framework (Höhna et al. 2014; Höhna et al. 2016), allows for easy implementation of the stepwise approach for, in principle, any hierarchical phylogenetic model. For instance, a joint model estimating gene trees, a species tree, and trait-dependent diversification rates (*e*.*g*., the BiSSE model (Maddison et al. 2007)) could be split into three steps in `RevBayes`, where the first step infers a distribution of gene trees, the second infers distributions for the species tree using an importance sample of those gene trees, and the third infers speciation and extinction rates using an importance sample of the rate parameters.

## Results

### Joint Bayesian inference vs. stepwise Bayesian inference

First, we validated our theory and implementation of stepwise Bayesian inference. We used a simple toy example where multiple sets of data *x*_*ij*_ were drawn from a Poisson distribution with parameter *λ*_*i*_. Each *λ*_*i*_ was drawn from a lognormal distribution with parameters *μ* and *σ*. Thus, our likelihood function was a composite Poisson distribution with parameters *λ*_*i*_, our prior distribution was a lognormal distribution with parameters *μ* and *σ*, and our hyper-prior distribution was a uniform distribution on *μ* and another uniform distribution on *σ* (see Figure 2). This toy example was motivated by the phylogenetic relaxed clock model where branch rates are drawn from a lognormal distribution and the number of substitutions on a branch follows a Poisson process.

Our goal was to determine whether the joint Bayesian inference and stepwise Bayesian inference approaches produced the same posterior distributions, as would be expected. The joint inference estimated the posterior distribution of our parameters *λ*_*i*_ and hyper-parameters *μ* and *σ* jointly. The stepwise inference first estimated an importance distribution for *λ*_*i*_ (*step 1*) and then the posterior distribution of *μ* and *σ* using a sample from the importance distribution (*step 2* ; see Figure 2). More details about the inference settings are provided in the *Supplementary Information*.

We tested different numbers of observations, *M* = {1, 10, 100}, as well as different numbers of samples in the second step of the stepwise Bayesian inference, *N* = {1, 10, 100, 1000} (additional results for *M* = 1000 are shown in Supplementary Figure S1). The varying number of observations simulated cases where the amount of information in the data varied; that is, the more samples we had, the more information there was in the data. Thus, we expected that we would obtain more precise estimates of *λ*_*i*_ in *step 1* with increasing *M*, and consequently more precise estimates of *μ* and *σ* in *step 2*. This hypothesis was based on the expectation that smaller amounts of error would be propagated from the importance distribution with increasing *M*.

Joint and stepwise inferences produced identical posterior distributions of *μ, σ*, and *λ* when the number of observations was 100 or 1000 (Figure 3 and Figure S1). When we used only one observation per *λ*_*i*_, joint and stepwise inference estimates differed noticeably (Figure 3 left column). This was not surprising because in this situation we have as many parameters *λ*_*i*_ as we do observations. The joint inference utilized the shared hyperprior distribution to inform the posterior distribution on *λ*_*i*_, which was thus narrower than the importance distribution (Figure 3 left column, bottom row). In this situation it was the hyperprior distribution on *λ*_*i*_ that drove the posterior distribution, rather than the data and likelihood. In real empirical applications it would be questionable whether these estimates would be trustworthy, given that the data are overwhelmed by the (hyper-)prior distribution.

**Figure 3:**
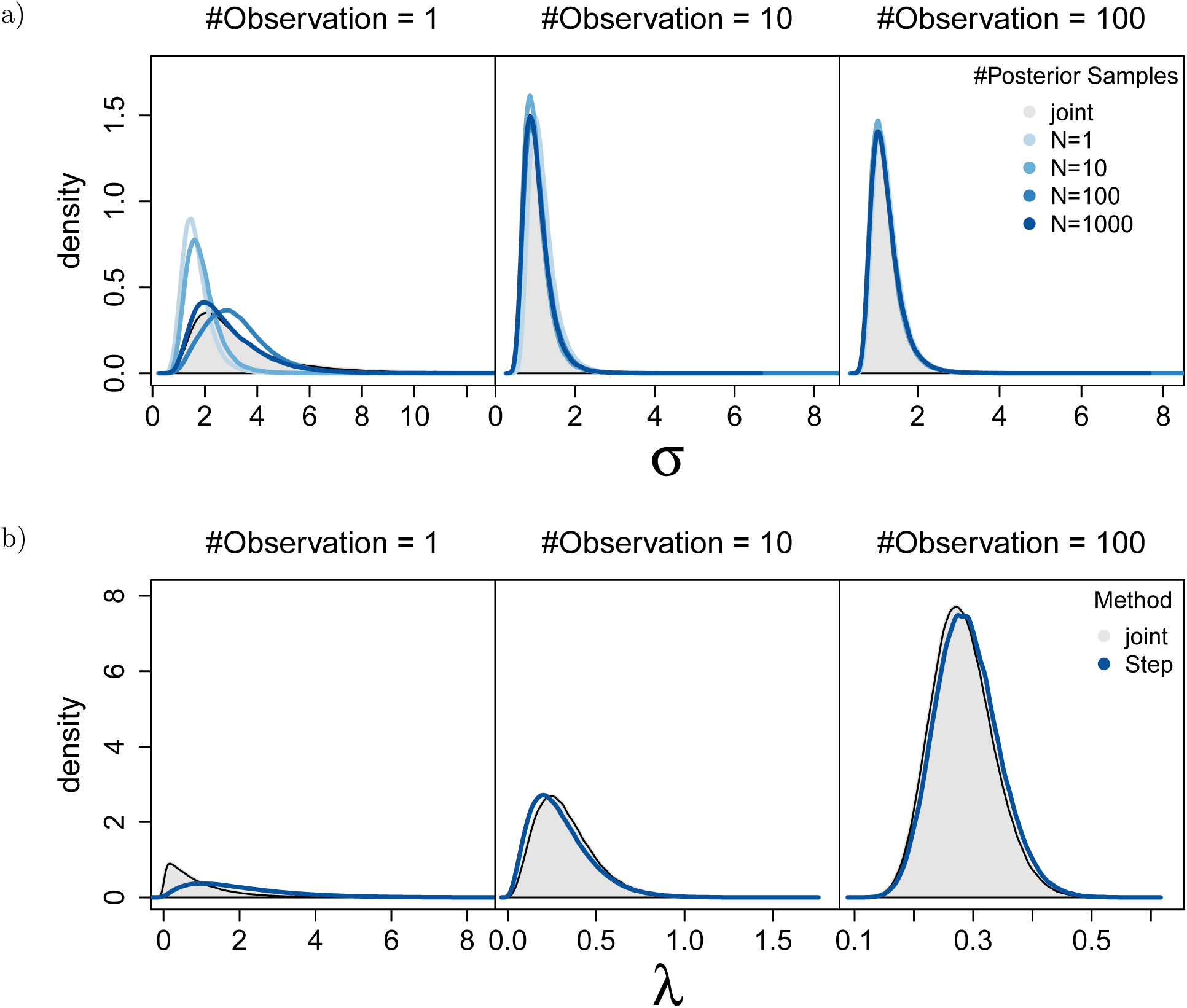
Estimated posterior distributions for the simple toy example as shown in Figure 2. In each row, we show the posterior distributions for different numbers of observations (columns) and different number of samples used in *step 2* (colors). The top row shows the posterior distributions of *σ*. The bottom row shows the posterior and importance distribution of an arbitrarily picked *λ*_*i*_. We observe that with few samples the joint and stepwise inferences disagree (left column; top row) which is due to divergence of the posterior distribution and importance distribution (left column; bottom row). Joint and stepwise inference are identical for many observations (right column). More samples *M* of the importance distribution in *step 2* are beneficial but have a smaller impact than the number of observations.

A larger number of samples *N* from the importance distribution should improve congruence between joint and stepwise inference. The number of samples *N* from the importance distribution did not impact the accuracy of the stepwise inference if the number of observations was large (Figure 3 right column). That is, when there was a high amount of information in the data, even a small number of samples from the importance distribution was sufficient to estimate the posterior with high accuracy. This robustness was due to the comparably narrow posterior distribution on *λ*_*i*_, which had virtually no outlier values. On the other side of the tested scenarios, where a small number of observations was used (Figure 3 left column), more samples from the importance distribution improved the congruence. In principle, if we had used infinitely many samples from the importance distribution, then even for the most difficult scenario — *i*.*e*., where we only have one observation per *λ*_*i*_ parameter — we would obtain identical results between joint and stepwise inference. Our results here indicate that this is in fact true and therefore validate our stepwise Bayesian inference theory and approach.

To develop an intuition for this observed behavior, imagine again that you want to compute the mean of a distribution. If there is (almost) no variation in the distribution, then even a single sample from the distribution will be very close to the mean value. Thus, one or very few samples would be sufficient. On the other hand, if the distribution has a large variance, then you would expect that any given sample would be further away from the mean than in a distribution with low variance. Thus, when variance is high, many more samples are necessary in order to estimate the distribution accurately.

### Divergence times estimation using stepwise Bayesian inference

The statistically most robust approach to estimating divergence times of a phylogeny from molecular sequence data is a Bayesian relaxed clock analysis (Thorne et al. 1998; Huelsenbeck et al. 2000a; Drummond et al. 2006). We refer readers to the large literature on divergence time estimation for more details (e.g., Donoghue and Yang 2016). The major complication in divergence time estimation is that the branch length *ν*_*i*_ is computed as the product of the branch rate *r*_*i*_ and the branch time *t*_*i*_, *i*.*e*., *ν*_*i*_ = *r*_*i*_ × *t*_*i*_. Thus, the branch length is not identifiable if neither the branch rate nor the branch time are known (Rannala 2016). This non-identifiability is resolved by somewhat informative prior distributions that give preference to more clock-like parameter settings (see Figure 4 in Donoghue and Yang (2016)).

**Figure 4:**
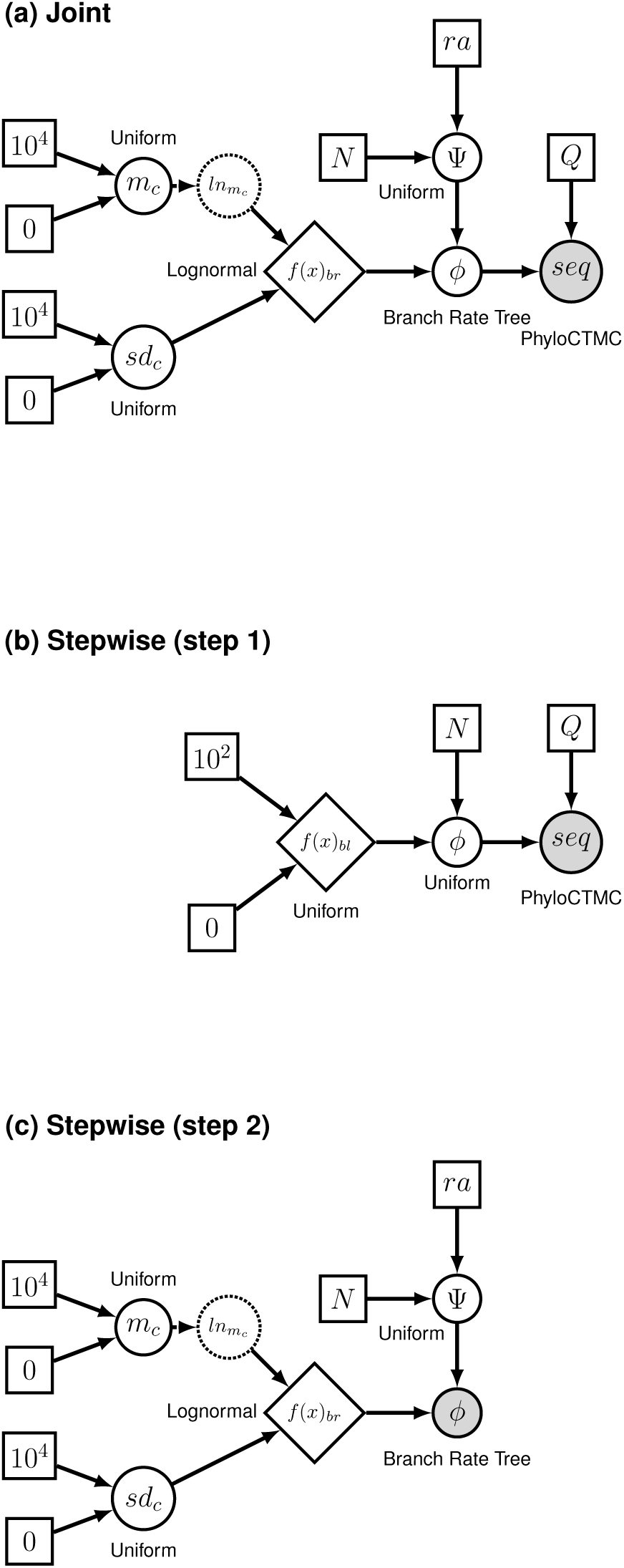
Graphical model of the joint (a) and stepwise (b,c) Bayesian inferences for the phylogenetic relaxed clock example. The rate matrix *Q* is unparametric Jukes-Cantor (Jukes and Cantor 1969) equal rates substitution matrix. In the joint model (a), the time tree Ψ is pulled from a uniform prior given the taxa *N* and the root age *ra*. Then, in combination with a lognormal distribution on the branch rates (*f* (*x*)_*br*_; note here that we use a diamond to represent a distribution node in a directed acyclic graph) with hyperparameters *m*_*c*_ and *sd*_*c*_, the distribution of unrooted branch length trees *ϕ* is generated using the branch rate tree distribution (dnBranchRateTree; see main text). All parameter values are then inferred using the sequence data (*seq*) via the standard phylogenetic continuous-time Markov chain (PhyloCTMC). In the stepwise approach, we first infer the posterior of branch length trees *ϕ* in *step 1* (b) using a uniform prior on the tree topology with taxa *N* and a uniform distribution over the interval [0, 100] on the branch lengths (*f* (*x*)_*bl*_). Then, in *step 2* (c), we use an importance sample of the posterior distribution of *ϕ* as data, and infer the divergence times using a uniform distribution on the time trees Ψ and the branch rate tree distribution.

MCMC analyses of these weakly identifiable hierarchical models are intrinsically hard and often fail to converge. Alternatively, the phylogeny can be estimated with branch lengths in units of substitutions per site first, with divergence times then estimated using one of several available adhoc methods (Sanderson 2003; Britton et al. 2007; Smith, Stephen A. and O’Meara, Brian C. 2012; Tamura et al. 2012; Rambaut et al. 2016; To et al. 2016). These ad-hoc stepwise approaches can perform well under certain circumstances (*e*.*g*., when there is a moderate amount of among branch rate variation), but are neither equivalent to nor as good as full Bayesian relaxed clock approaches (Tong et al. 2018).

Here we develop and test a stepwise Bayesian relaxed clock approach for divergence time estimation that is equivalent to joint inference of branch lengths, branch rates, and divergence times (Figure 4). The stepwise Bayesian relaxed clock analysis consists of the following two steps: (1) estimation of the posterior distribution of unrooted phylogenies (tree topologies and branch lengths), and (2) estimation of the posterior distribution of divergence times using an importance sample from the posterior inferred in step (1).

Traditionally, joint Bayesian inference of divergence times and branch-specific molecular clock rates uses a prior distribution on the branch rates and another prior distribution on the timetree (*i*.*e*., the ultrametric phylogeny with divergence times). Then, the branch times are multiplied by the branch rates to obtain the branch lengths when computing the likelihood of the observed molecular sequence data (Höhna et al. 2016). That is, the model consists of a stochastic timetree variable Ψ and a vector of stochastic branch rate variables **r**. There is no variable that corresponds to an unrooted phylogeny with branch lengths, leading to problems for the stepwise Bayesian analysis because, as formulated, the result of the first step should be a posterior distribution of unrooted phylogenies with branch lengths. Therefore, we restructure the traditional relaxed clock model to use a *branch-rate-tree distribution*. The details of this new distribution are provided in the *Material and Methods*. Interestingly, our new branch-rate-tree distribution shows a philosophically different view on the tree variable, although the overall model is identical to traditional relaxed clock models (Figure 4). Namely, in our model, a tree with branch lengths is the intrinsical parameter of a model of molecular character data evolution, rather than an ultrametric tree with divergence times together with a vector of branch rates.

First, we simulated phylogenetic trees under a birth-death process with speciation rate *b* = 1.0 and extinction rate *d* = 0.5 conditioned on observing *T* = {8, 16, 32} taxa using the R package `TESS` (Höhna 2013; Höhna et al. 2016). Afterwards we rescaled the tree to a root age of 5.0. Second, we simulated sequence alignments with *K* = {100, 1000, 10000} sites under a Jukes-Cantor substitution model (Jukes and Cantor 1969) and a lognormal relaxed clock model with mean *μ* = 0.1 and standard deviation *σ* = 0.3. We kept the substitution model purposefully simple to keep the computational time of the analyses low. Then, we analyzed each simulated dataset under a joint Bayesian analysis and five stepwise Bayesian relaxed clock analyses where the number of samples from the posterior distribution of branch-rate trees was *N* = {1, 10, 100, 1000}. The details of the MCMC simulations are provided in the *Supplementary Information*.

We observed that the estimated posterior distributions of the parameters *μ* and *σ* of the lognormal prior distribution on the branch-specific clock rates were identical between the joint Bayesian relaxed clock analysis and the stepwise Bayesian relaxed clock analysis if the number of sites was large (Figure 5a). This result confirmed our conclusion from the small toy example that the step-wise Bayesian analysis resembled the joint Bayesian analysis more closely if the information in the data was high. In this case, even as few as *N* = 10 samples from the posterior distribution in *step 1* could be sufficient. If the observations were intermediate in number (1000 sites), then at least *N* = 100 samples from the posterior distribution of branch-rate trees were needed to obtain a very good agreement between the two approaches. If the information in the data was low (100 sites), then even *N* = 1000 samples were not sufficient to provide a close enough approximation of the posterior distribution. These results show the expected behavior: datasets with low information have wider posterior distributions (and thus more samples are needed for a good approximation), and the impact of the prior is stronger (reflecting the difference in hierarchical priors of joint vs. stepwise inferences).

**Figure 5:**
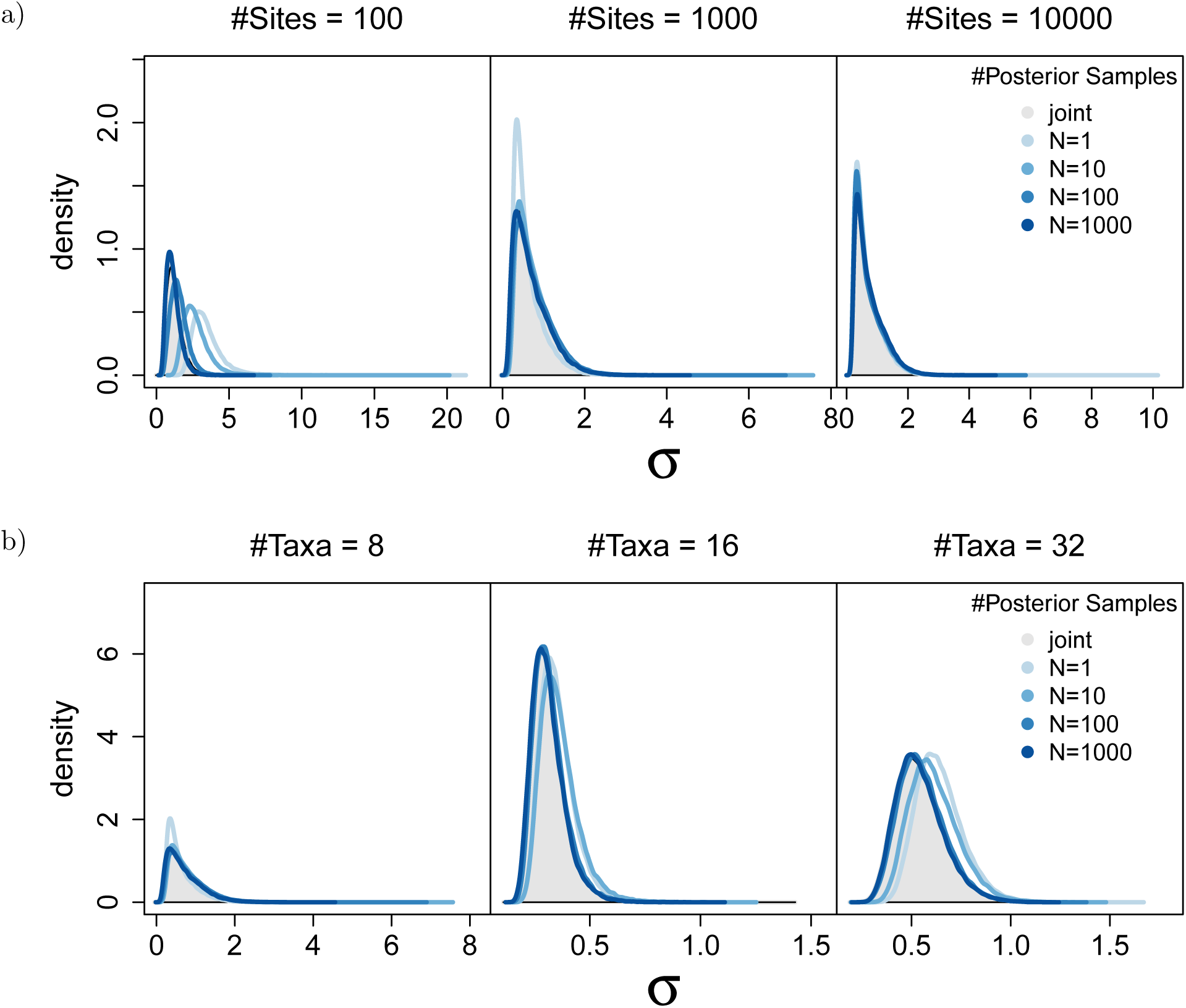
Estimated posterior distributions of the hyper-parameter *σ*, the standard deviation of the log-normally distributed branch rates, for the model shown in Figure 4. The posterior distribution of *σ* for the joint Bayesian analysis is shown in gray and in different colors (light to dark blue) for increasing number of samples in the second step of the stepwise Bayesian analysis. The three plots in the top row (a) show one example each for a different number of simulated sites: 100, 1000 and 10000 (from left to right). The number of taxa was 8 for these analyses. As with the toy example (Figure 3) we observe that both more data (*i*.*e*., number of sites) and more samples *N* of the posterior distribution in the stepwise Bayesian approach improve the accuracy. The three plots in the bottom row (b) show one example each for a different number of taxa: 8, 16 and 32 (from left to right). The number of sites for these analyses was 1000. Here, more taxa appear to produce more discrepancy between the joint and stepwise Bayesian analysis because fewer information per branch is contained in the same number of sites (lower sites to branches ratio). However, the scale of these posterior distribution changes because more branches lead to more information on the branch-rate hyperprior parameters. Other aspects of the simulation and analyses are described in the main text.

Varying the number of taxa showed results that were more difficult to interpret (Figure 5b). If a dataset contained more taxa, but the same number of sites, then the ratio of sites to branches was lower. As a result, there would be less information for each branch, leading to higher uncertainty per branch. However, when there were more branches, there was more information about the variation in branch-specific clock rates, which in turn decreased the uncertainty in the posterior distribution of the lognormal branch rate prior (as evidenced by the narrower posterior distribution in Figure 5b). This trade-off resulted in the more complex behavior shown in Figure 5b.

From a biological perspective, the focal parameters of relaxed clock analyses are the divergence times (or node ages). We therefore compared the node age estimates of the joint Bayesian relaxed clock analyses to the node age estimates of the stepwise Bayesian relaxed clock analyses (Figure 6). In agreement with our previous results, the age estimates of the two different approaches were identical if the information in the data was high (*e*.*g*., 10,000 sites). If the number of sites was intermediate (1,000 sites), then more samples (e.g., *N* = 100) from the posterior distribution of branch-rate trees were needed for congruence. Furthermore, we observed that the sites-to-branches ratio has a strong impact on our ability to estimate node ages (that is, more sites resulted in improved estimates, but more branches resulted in less accurate estimates).

**Figure 6:**
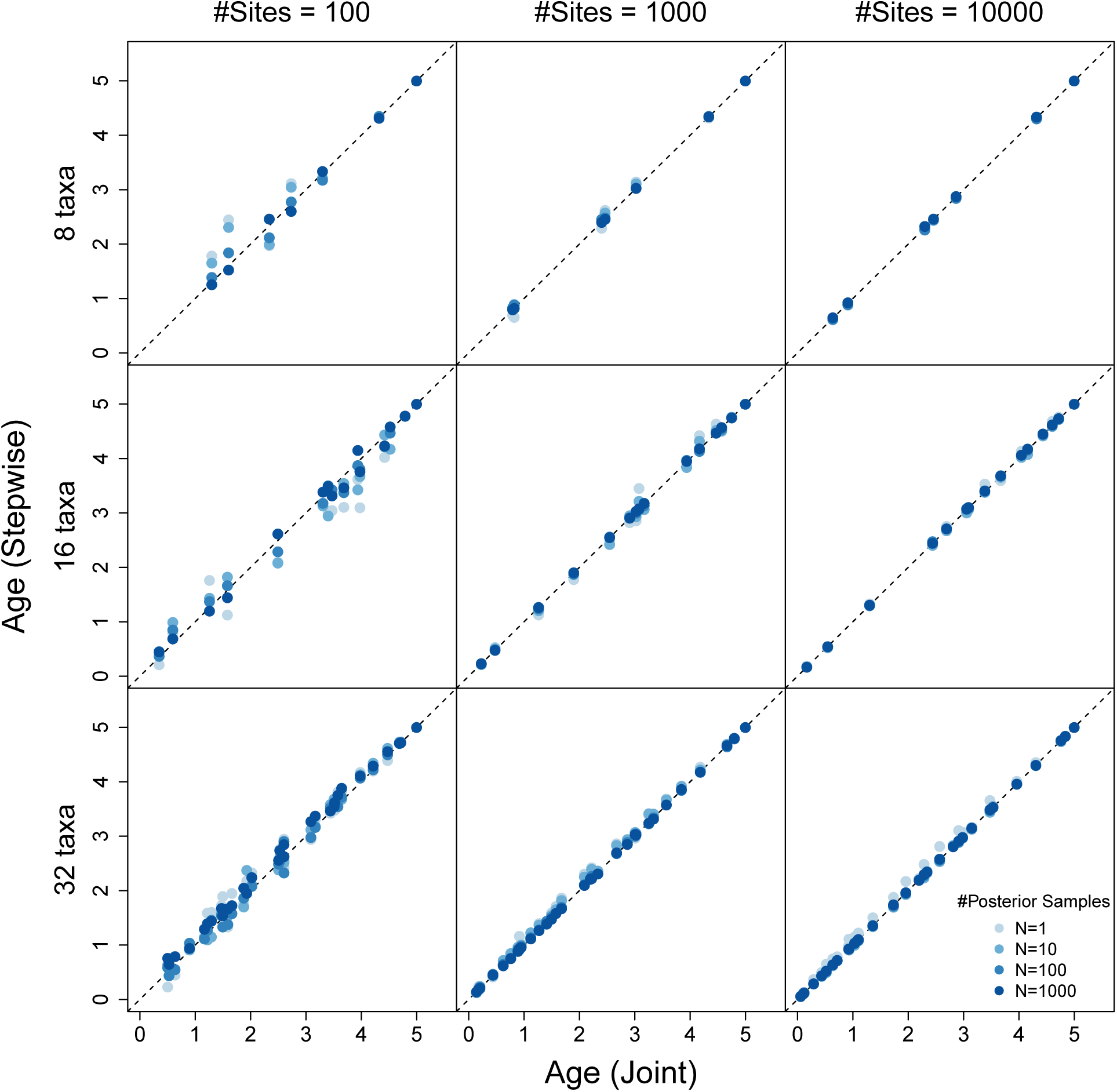
Estimated posterior distributions of node ages for the relaxed clock analyses as shown in Figure 4. The mean posterior estimate of the node ages for the joint Bayesian relaxed clock analysis is shown on the x-axis and the mean posterior estimate of the node ages for the stepwise Bayesian relaxed clock analysis are shown on the y-axis, in different colors (green to blue) for increasing number of samples in the second step of the stepwise Bayesian analysis. The three plots show one example each for a different number of sites: 100, 1000 and 10000 (from left to right). The three columns show different number of taxa: 8, 16 and 32 (from top to bottom). The agreement between the two approaches increases with more sites but decreases with more taxa.

## Discussion

### Number of samples

Most existing ad-hoc two-step methods, such as `RelTime` (Tamura et al. 2012, 2018), `pathd8` (Britton et al. 2007), `treePL` (Smith, Stephen A. and O’Meara, Brian C. 2012), `LSD` (To et al. 2016), and `TempEst` (Rambaut et al. 2016), use only a single estimated phylogeny with branch lengths as input data to estimate divergence times. The program `r8s` (Sanderson 2003) can take multiple trees with different branch lengths as input to calculate confidence intervals on the estimated ages, but these trees must all share the same topology. Thus, this method does not account for uncertainty in the tree topology during divergence time estimation. In theory, a large number of samples are necessary in order to obtain a good approximation of the branch length posteriors and agreement between joint and stepwise Bayesian analyses. Hence, a researcher is left with the question: “How many samples are necessary for a good approximation?”

Our results are mixed. In some situations, a single sample might be sufficient, whereas in other situations, many samples (*e*.*g*., *N* ≥ 1000) might be necessary (Figures 5–6). To be on the safe side, we recommend to use as many samples as possible (at least *N* ≥ 100). However, in practice the number of samples is necessarily limited by the concomitant increase in computational cost (see discussion below).

Another possible solution is to take several sets of samples of a small or intermediate amount, *e*.*g*., *N* = 10 or *N* = 100. Then, the second step of the stepwise Bayesian analysis is repeated and the difference between the resulting posterior distributions is computed using a metric such as the Kolmogorov-Smirnov test. The number of samples can be deemed to be sufficient when the posterior distributions resulting from different sample sets are acceptably similar (which would depend on the metric used; Figure 7). The increased computational time required to run these repeated tests is mitigated by the fact that the posterior from the first step (during which the computationally intensive calculation of the likelihood of the data is performed) needs only to be estimated once.

**Figure 7:**
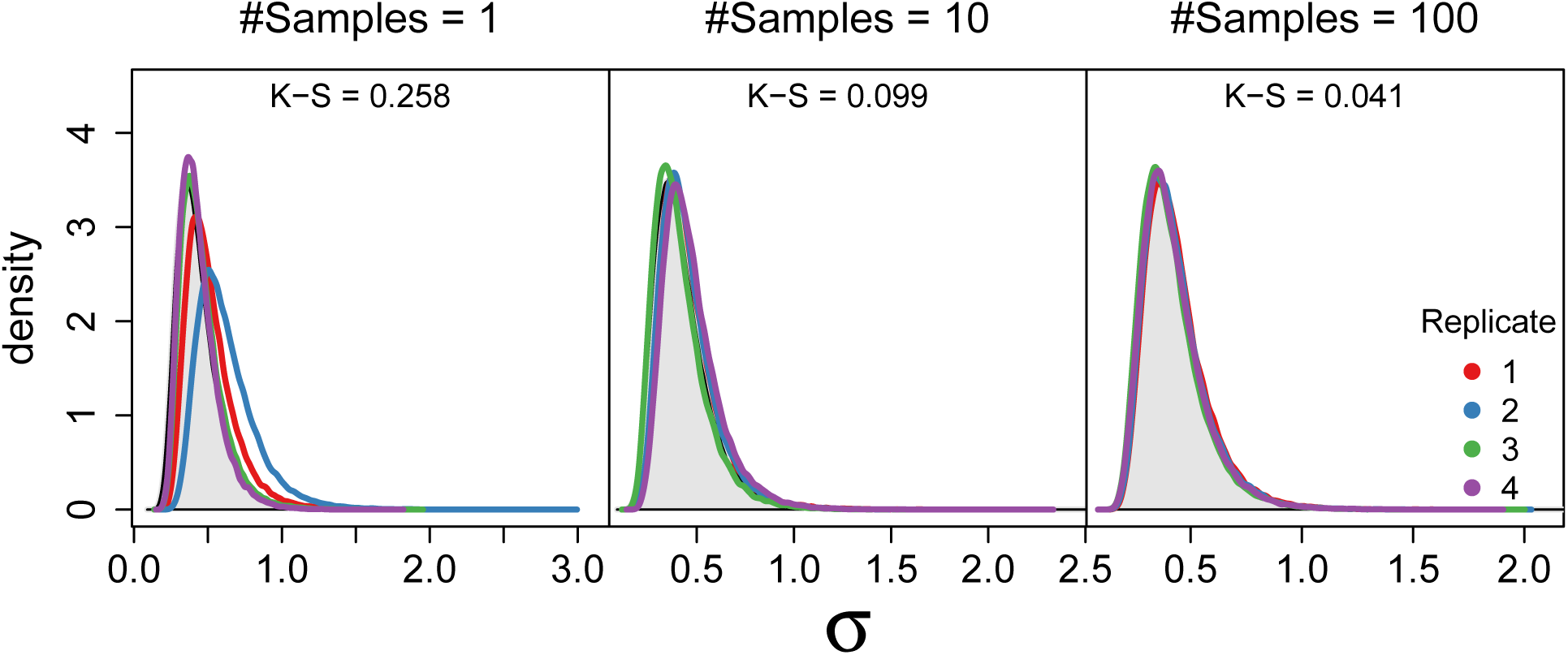
Posterior distributions of *σ* computed using the stepwise Bayesian relaxed clock model. The shaded gray distribution represents the posterior distribution of the joint Bayesian relaxed clock model as a reference. The colored solid lines represent different repetitions with different samples taken from the distribution of branch-length-trees from step 1. We used *N* = {1, 10, 100} samples in step 2 (from left to right). For small *N* (left panel) we observe that different repetitions yield different posterior distributions. This difference in posterior distribution is significant by the average Kolmogorov-Smirnov (K-S) score.

### Computation speed

The stepwise Bayesian inference can be computationally more efficient than traditional joint Bayesian inferences (Figure 8). For example, the joint inference has an ESS per second of 4.5 for our simulated dataset with 1000 sites. For the same dataset, *step 1* has an ESS per second of 7.4 and *step 2* with 100 samples from the posterior distribution has an ESS per second of 9.2. Thus, the joint analysis needs 44.4 seconds to reach 200 independent samples while both *step 1* and *step 2* together need 48.8 seconds. In our example, both approaches are approximately equally computationally efficient, although it can be seen that the burden of the stepwise Bayesian inference lies in *step 1*, which includes expensive computation of the probability of the data.

**Figure 8:**
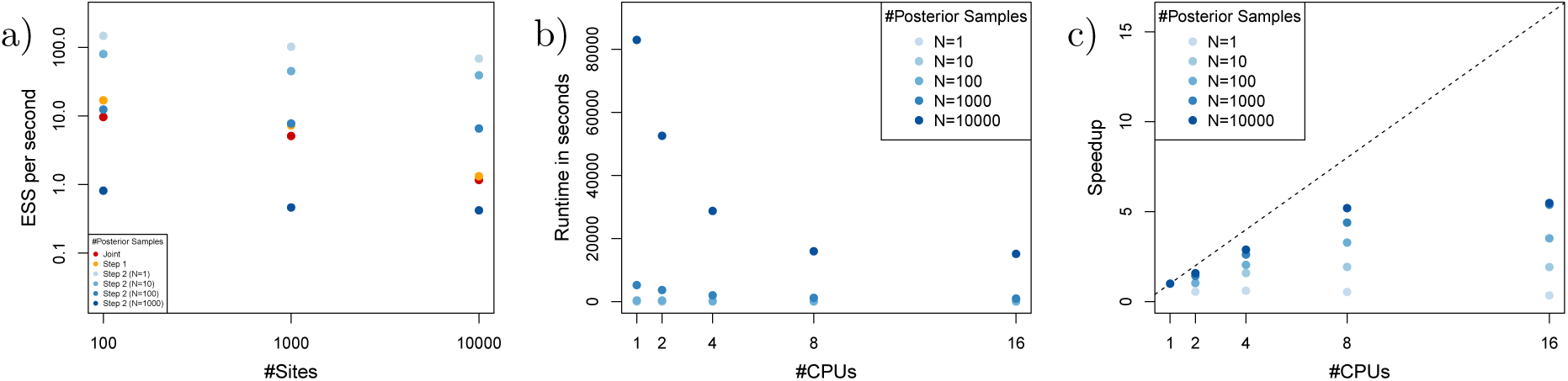
Performance results comparing the runtime and sampling efficiency of the joint and stepwise Bayesian approach. a) The average effective sample size (ESS) per second for each type of analysis. b) Average runtime of the second step of the stepwise Bayesian relaxed clock analysis with different number of samples and different number of CPUs. c) Speedup for step 2 when using multiple CPUs.

Furthermore, to test different models, *e*.*g*., different relaxed clock models, one only needs to redo *step 2*. Since *step 2* with 1–100 samples from the posterior distribution of *step 1* is more efficient in terms of ESS per second, it is faster with the stepwise Bayesian inference to perform these analyses.

Finally, *step 2* can be easily parallelized. The central equation of the likelihood in *step 2* involves the sum over all samples (Equation 7). If the number of samples is sufficiently large, then it makes sense to distribute the individual likelihood computations within the sum over many processors (*e*.*g*., ten times as many samples as processors). Thus, *step 2* can be performed efficiently on common HPC infrastructures (Figure 8b). We note that we have not optimized our parallel code which still shows some unnecessary overhead but it works sufficiently well to illustrate the potential of using many samples while computing the probabilities in parallel.

## Conclusions

The stepwise approach we describe here can be applied to any sequence of Bayesian analyses. Doing so allows for uncertainty to be properly incorporated and propagated through a bioinformatics pipeline, resulting in more robust and accurate inferences. Disregarding uncertainty, *i*.*e*., taking only a single point estimate, in the next step of a bioinformatics pipeline can lead to biased estimates (Figures 5 and 6)

While some variant of the stepwise approach has been applied to phylogenetic questions before (*e*.*g*., only for combined data analyses instead of complex hierarchical models), we present here the first formal demonstration and explicit testing of the equivalence between the joint and stepwise approaches. Further, our method represents the first application of the full Bayesian stepwise approach to divergence time estimation. We have provided the theoretical formulation of the stepwise approach and shown that it can be applied to hierarchical models of any arbitrary number of layers. While we only present the results from a hierarchical model of two layers here, in principle any joint model can be decomposed into a series of stepwise analyses. Computationally, this potentially allows for complex, parameter-rich models that are computationally intractable using a joint approach to be discretized into several smaller, computationally feasible steps. In this case, for example, the second step dispenses with the expensive calculation of the likelihood of the sequence data, resulting in very fast analyses (see Figure 8a). As we have shown, this stepwise approach would be equivalent to the joint analysis. The stepwise approach thus not only allows for the vital incorporation of uncertainty in bioinformatics pipelines, but also potentially opens up the types and scales of analyses that are possible given current computational capabilities.

Finally, with this contribution to the theory of bioinformatics pipelines we hope that both method developers and applied researchers will consider the stepwise Bayesian inference framework to propagate uncertainty. That is, new methods need to be able to use not only a single dataset but a sample of datasets as input and results need to be archived as samples instead of point estimates.

## Acknowledgements

We thank Bastien Boussau and Gergely Szöllősi for discussions stepwise inference that have lead to this research. We also want to thank Jiansi Gao for his insights into the stepwise Bayesian approach employed in BEAST. This research was supported by the Deutsche Forschungsgemeinschaft (DFG) Emmy Noether-Program HO 6201/1-1 awarded to SH.

## Notes

### Competing Interest Statement

The authors have declared no competing interest.

## References

Britton, T., C. L. Anderson, D. Jacquet, S. Lundqvist, and K. Bremer. 2007. Estimating Divergence Times in Large Phylogenetic Trees. Systematic Biology 56:741–752.

Donoghue, P. C. J. and Z. Yang. 2016. The evolution of methods for establishing evolutionary timescales. Philosophical Transactions of the Royal Society B: Biological Sciences 371:20160020.

Drummond, A. J., S. Y. W. Ho, M. J. Phillips, and A. Rambaut. 2006. Relaxed Phylogenetics and Dating with Confidence. PLoS Biology 4:e88.

Faria, N.R., R. A. M. A. Suchard, G. Baele, T. Bedford, M. J. Ward, A. J. Tatem, J. D. Sousa, N. Arinaminpathy, J. Pépin, D. Posada, M. Peeters, O. G. Pybus, and P. Lemey. 2014. The early spread and epidemic ignition of HIV-1 in human populations. Science 56:56–61.

Hastings, W. K. 1970. Monte Carlo Sampling Methods Using Markov Chains and Their Applications. Biometrika 57:97–109.

Höhna, S. 2013. Fast simulation of reconstructed phylogenies under global time-dependent birth-death processes. Bioinformatics 29:1367–1374.

Höhna, S., T. A. Heath, B. Boussau, M. J. Landis, F. Ronquist, and J. P. Huelsenbeck. 2014. Probabilistic Graphical Model Representation in Phylogenetics. Systematic Biology 63:753–771.

Höhna, S., M. J. Landis, T. A. Heath, B. Boussau, N. Lartillot, B. R. Moore, J. P. Huelsenbeck, and F. Ronquist. 2016. RevBayes: Bayesian phylogenetic inference using graphical models and an interactive model-specification language. Systematic Biology 65:726–736.

Höhna, S., M. R. May, and B. R. Moore. 2016. TESS: an R package for efficiently simulating phylogenetic trees and performing Bayesian inference of lineage diversification rates. Bioinformatics 32:789–791.

Huelsenbeck, J. P., B. Larget, and D. L. Swofford. 2000a. A compound Poisson process for relaxing the molecular clock. Genetics 154:1879–1892.

Huelsenbeck, J. P. and B. Rannala. 2003. Detecting correlation between characters in a comparative analysis with uncertain phylogeny. Evolution 57:1237–1247.

Huelsenbeck, J. P., B. Rannala, and J. P. Masly. 2000b. Accommodating phylogenetic uncertainty in evolutionary studies. Science 288:2349–2350.

Jukes, T. and C. Cantor. 1969. Evolution of protein molecules. Mammalian Protein Metabolism 3:21–132.

Maddison, W., P. Midford, and S. Otto. 2007. Estimating a binary character’s effect on speciation and extinction. Systematic Biology 56:701–710.

Metropolis, N., A. W. Rosenbluth, M. N. Rosenbluth, A. H. Teller, and E. Teller. 1953. Equation of State Calculations by Fast Computing Machines. Journal of Chemical Physics 21:1087–1092.

Nylander, J. A. A., U. Olsson, P. Alström, and I. Sanmartín. 2008. Accounting for phylogenetic uncertainty in biogeography: a Bayesian approach to dispersal-vicariance analysis of the thrushes (Aves: Turdus). Systematic Biology 57:257–268.

Pagel, M. and F. Lutzoni. 2002. Biological Evolution and Statistical Physics chap. Accounting for phylogenetic uncertainty in comparative studies of evolution and adaptation, Pages 148–161. Springer.

Pagel, M., A. Meade, and D. Barker. 2004. Bayesian estimation of ancestral character states on phylogenies. Systematic Biology 53:673–684.

Rambaut, A., T. T. Lam, L. Max Carvalho, and O. G. Pybus. 2016. Exploring the temporal structure of heterochronous sequences using TempEst (formerly Path-O-Gen). Virus evolution 2:vew007.

Rannala, B. 2016. Conceptual issues in Bayesian divergence time estimation. Philosophical Transactions of the Royal Society B: Biological Sciences 371:20150134.

Sanderson, M. J. 2003. r8s: inferring absolute rates of molecular evolution and divergence times in the absence of a molecular clock. Bioinformatics 19:301–302.

Smith, Stephen A. and O’Meara Brian C. 2012. treePL: divergence time estimation using penalized likelihood for large phylogenies. Bioinformatics 28:2689–2690.

Tamura, K., F. U. Battistuzzi, P. Billing-Ross, O. Murillo, A. Filipski, and S. Kumar. 2012. Estimating divergence times in large molecular phylogenies. Proceedings of the National Academy of Sciences 109:19333–19338.

Tamura, K., Q. Tao, and S. Kumar. 2018. Theoretical foundation of the reltime method for estimating divergence times from variable evolutionary rates. Molecular Biology and Evolution 35:1770–1782.

Thorne, J., H. Kishino, and I. S. Painter. 1998. Estimating the rate of evolution of the rate of molecular evolution. Molecular Biology and Evolution 15:1647–1657.

To, T.-H., M. Jung, S. Lycett, and O. Gascuel. 2016. Fast dating using least-squares criteria and algorithms. Systematic Biology 65:82–97.

Tong, K. J., D. A. Duchëne, S. Duchëne, J. L. Geoghegan, and S. Y. W. Ho. 2018. A comparison of methods for estimating substitution rates from ancient DNA sequence data. BMC Evolutionary Biology 18:70.

