## Supplementary Material for "Stepwise Bayesian Phylogenetic Inference"

#### Supporting Information

#### Contents


S1 Toy example . . . . . . . . . . . . . . . . . . . . . . . . . . . . . . . . . . . . . . . . . . S2
S1.1 Joint inference . . . . . . . . . . . . . . . . . . . . . . . . . . . . . . . . . . . S2
S1.2 Stepwise Bayesian inference . . . . . . . . . . . . . . . . . . . . . . . . . . . S3
S1.2.1 Step 1 . . . . . . . . . . . . . . . . . . . . . . . . . . . . . . . . . . . . S3
S1.2.2 Step 2 . . . . . . . . . . . . . . . . . . . . . . . . . . . . . . . . . . . . S4
S2 Relaxed Clock Analyses . . . . . . . . . . . . . . . . . . . . . . . . . . . . . . . . . . . S5
S2.1 Joint inference . . . . . . . . . . . . . . . . . . . . . . . . . . . . . . . . . . . S5
S2.2 Stepwise Bayesian inference . . . . . . . . . . . . . . . . . . . . . . . . . . . S6
S2.2.1 Step 1 . . . . . . . . . . . . . . . . . . . . . . . . . . . . . . . . . . . . S6
S2.2.2 Step 2 . . . . . . . . . . . . . . . . . . . . . . . . . . . . . . . . . . . . S7
S2.3 Results . . . . . . . . . . . . . . . . . . . . . . . . . . . . . . . . . . . . . . . S8

### S1 Toy example

In this section we provide the details for the Markov chain Monte Carlo (MCMC) analyses on the toy example (see Figure 1 in the main text).

#### S1.1 Joint inference

The parameters and prior distribution for the joint inference of toy example corresponding to Figure 1a are given in Table S1. A corresponding RevBayes script outlining all necessary details for replicating this analysis are given in Listing 1. We ran the MCMC analyses for 1,000,000 iterations, after a pre-burnin phase of 100,000 iterations, which is well beyond the normal chain length but the obtain very smooth posterior distributions. The specific moves on the parameters can be retrieved from the listing.

**Table S1: Model parameter names and prior distributions for the toy example joint inference.**

| Parameter | $X$ | $f(X)$ |
|---|---|---|
| Prior mean | $\mu$ | Uniform$(-10, 10)$ |
| Prior standard deviation | $\sigma$ | Uniform$(0, 10^4)$ |
| Focal parameter | $\lambda_i$ | Lognormal$(\mu, \sigma)$ |
| Observations | $x_{i,j}$ | Poisson$(\lambda_i)$ |

```
mu ~ dnUniform(-10,10)
sigma ~ dnUniform(0,1E4)

idx <- 1
for (i in 1:10) {
  lambdas[i] ~ dnLognormal(mu, sigma)
  for (j in 1:M ) {
    x[idx] ~ dnPoisson(lambdas[i])
    x[idx].clamp(data[idx])
    idx++
  }
}

moves.append( mvSlide(mu, delta=0.01) )
moves.append( mvScale(sigma, lambda=0.01) )

for (i in 1:NUM_LAMBDAS) {
  moves.append( mvScale(lambdas[i], lambda=0.01) )
}

monitors.append( mnModel(filename="output/joint_rep.log", printgen=10) )
monitors.append( mnScreen(mu, sigma, printgen=1000) )

mymcmc = mcmc(mymodel, monitors, moves)
```

```
mymcmc.burnin(1E5,100)
mymcmc.run(1E6)
```

**Listing 1:** Excerpt from the RevBayes script for the joint inference on the toy example.

#### S1.2 Stepwise Bayesian inference

In this next subsection, we provide more details about the stepwise Bayesian inference for the toy example.

##### S1.2.1 Step 1

*Step 1* of the stepwise Bayesian inference estimates the importance distribution of the focal parameter. Thus, the hierarchical layer of the model is broken up and only the data layers are included (see Table S2 and Figure 1b). Overall, *step 1* is very similar to the joint inference with the exception that the hierarchical prior distribution is replaced by a uniform prior distribution. The remaining MCMC settings stayed the same. Listing 2 shows the RevBayes script to perform *step 1* of the stepwise Bayesian inference.

**Table S2: Model parameter names and prior distributions for *step 1* of the stepwise Bayesian inference for the toy example.**

| Parameter | $X$ | $f(X)$ |
|---|---|---|
| Focal parameter | $\lambda_i$ | Uniform$(0, 10^4)$ |
| Observations | $x_{i,j}$ | Poisson$(\lambda_i)$ |

```
for (i in 1:10) {
  lambdas[i] ~ dnUniform(0,1E4)
  for (j in 1:M) {
    x[idx] ~ dnPoisson(lambdas[i])
    x[idx].clamp(data[idx])
    idx++
  }
}

for (i in 1:10) {
  moves.append( mvScale(lambdas[i], lambda=0.01) )
}

monitors.append( mnFile(lambdas, filename="output/step1.log", printgen=10) )
monitors.append( mnScreen(printgen=1000) )

mymcmc = mcmc(mymodel, monitors, moves)
mymcmc.burnin(1E5,100)
mymcmc.run(1E6)
```

**Listing 2:** Excerpt from the RevBayes script for *Step 1* of the stepwise Bayesian inference on the toy example.

##### S1.2.2 Step 2

**Table S3: Model parameter names and prior distributions for *step 2* of the stepwise Bayesian inference for the toy example.**

| Parameter | $X$ | $f(X)$ |
|---|---|---|
| Prior mean | $\mu$ | Uniform$(-10, 10)$ |
| Prior standard deviation | $\sigma$ | Uniform$(0, 10^4)$ |
| Focal parameter | $\lambda_i$ | Lognormal$(\mu, \sigma)$ |

*Step 2* of the Bayesian stepwise inference takes the samples from the importance distribution generated in *step 1* as data. We achieve this using the function `readTrace`. How many samples are actually taken is controlled by the thinning argument, that is, when we performed our tests using a different number of samples $N$ from *step 1* we simply applied a different thinning of the samples. The parameters and distributions for *step 2* are shown in Table S3 (see also Figure 1c). Compared to the joint inference, we do not have the layer including the data. Note that we use `dnEmpiricalSample` to represent the probability distribution for the importance sample. This `dnEmpiricalSample` is completely generic and can be used for any probability distribution.

```
samples = readTrace(file="output/step1.log", burnin=0.001, thin=1000)

mu ~ dnUniform(-10,10)
sigma ~ dnUniform(0,1E4)

for (i in 1:10) {
   lambda_prior[i] = dnLognormal(mu, sigma)
   lambdas[i] ~ dnEmpiricalSample(lambda_prior[i])
   lambdas[i].clamp( samples[4+i].getValues() )
}

moves.append( mvScale(sigma, lambda=0.01) )
moves.append( mvSlide(mu, delta=0.01) )

monitors.append( mnFile(mu, sigma, filename="output/step2.log", printgen=10) )
mymcmc.burnin(1E5,100)
mymcmc.run(1E6)
```

**Listing 3:** Excerpt from the RevBayes script for *Step 2* of the stepwise Bayesian inference on the toy example.

### S2 Relaxed Clock Analyses

In this section we describe the details of the relaxed clock analyses of the main text. The RevBayes code snippets describe the main features of the analyses and only need to be adapted for the specific data (*e.g.,* simulation replicate).

#### S2.1 Joint inference

First, we present the model for the joint Bayesian inference using the uncorrelated lognormal relaxed clock model. All parameters are presented in Table S4. We used a Jukes-Cantor substitution model [1] for simplicity and to speed up the simulation study. We used a uniform prior on node ages conditional on the root age. We used a lognormal prior distribution on branch-specific clock rates (uncorrelated lognormal relaxed clock [2]) with mean $m_c$ (in real space, *i.e.,* not log-transformed) and standard deviation $sd_c$. Both hyperparameter had a uniform prior distribution between 0 and 10,000.

**Table S4: Model parameter names and prior distributions for the Bayesian relaxed clock example joint inference, see main text Figure 3.**

| Parameter | $X$ | $f(X)$ |
|---|---|---|
| Substitution rate matrix | $Q$ | fixed to Jukes-Cantor model [1] |
| Mean clock rate | $m_c$ | $\text{Uniform}(0, 10^4)$ |
| Standard deviation of clock rates | $sd_c$ | $\text{Uniform}(0, 10^4)$ |
| Phylogeny | $\Psi$ | UniformTimeTree(*root*) |

The corresponding RevBayes code snippet is given in Listing 4. Note that we use our newly developed distribution `dnBranchRateTree` here as the prior distribution on the phylogeny with branch lengths in units of substitutions. In principle, the `root_branch_fraction` should not be a parameter but an observation and thus should not have a prior distribution. Here we used a uniform prior distribution that does not affect the outcome because all values are equally probable. Ideally, our implementation would integrate analytically or numerically over the `root_branch_fraction`. This might be implemented in a future version.

The MCMC simulation was run again for 1,000,000 iterations after a pre-burnin phase of 100,000 iterations. Note that multiple moves were applied per iteration where the average number of moves corresponds to the specified weights. That is, for 8 taxa we used 31 moves per iterations.

```
Q <- fnJC(4)

clock_rate_mean ~ dnUniform(0,1E4)
clock_rate_sd ~ dnUniform(0,1E4)

clock_rate_ln_mean := ln(clock_rate_mean)
branch_rate_prior = dnLognormal(clock_rate_ln_mean, clock_rate_sd)

time_tree ~ dnUniformTimeTree(rootAge=ROOT_AGE, taxa=taxa)
time_tree.setValue( true_tree )

```

```
root_branch_fraction ~ dnBeta(1,1)

psi ~ dnBranchRateTree( time_tree, branch_rate_prior, root_branch_fraction )

seq ~ dnPhyloCTMC(tree=psi, Q=Q, type="DNA")
seq.clamp(data)

moves.append( mvScale( clock_rate_mean, weight=3 ) )
moves.append( mvScale( clock_rate_sd, weight=3 ) )
moves.append( mvNodeTimeSlideUniform(time_tree,weight=NUM_TAXA) )
moves.append( mvBetaProbability(root_branch_fraction, weight=2.0) )
moves.append( mvBranchLengthScale(psi, weight=n_branches) )

monitors.append( mnFile(time_tree, filename="output/joint.trees", printgen=1) )
monitors.append( mnModel(filename="output/joint.log", printgen=1) )

mymcmc.burnin(1E5,100)
mymcmc.run(1E6)
```

**Listing 4:** Excerpt from the RevBayes script for the joint inference using the Bayesian relaxed clock example.

#### S2.2 Stepwise Bayesian inference

In this next subsection, we provide more details about the stepwise Bayesian inference for the Bayesian relaxed clock example.

##### S2.2.1 Step 1

*Step 1* of the stepwise Bayesian inference estimates the importance distribution of the phylogeny with branch lengths in units of subsitutions. Thus, the hierarchical layer of the model is broken up and only the data layers are included (see Table S2 and Figure 3b). Overall, *step 1* is very similar to the joint inference with the exception that the uniform node age prior distribution on the time tree together with the branch rate prior distribution is replaced by a prior distribution directly on unrooted trees (`dnUniformTopologyBranchLength`). The remaining MCMC settings stayed the same. Listing 5 shows the RevBayes script to perform *step 1* of the stepwise Bayesian inference.

**Table S5: Model parameter names and prior distributions for *step 1* of the stepwise Bayesian inference for the Bayesian relaxed clock example.**

| Parameter | $X$ | $f(X)$ |
|---|---|---|
| Substitution rate matrix | $Q$ | fixed to Jukes-Cantor model [1] |
| Phylogeny | $\Psi$ | UniformBranchLengthTree($blPrior = Uniform(0, 1000)$) |

```
Q <- fnJC(4)

psi ~ dnUniformTopologyBranchLength( taxa, dnUniform(0,1E3) )
```

```
psi.setValue( true_tree_unrooted )

moves.append( mvBranchLengthScale(psi, weight=n_branches) )

monitors.append( mnFile(psi, filename="output/step1.trees", printgen=10) )
monitors.append( mnScreen(printgen=1000) )

mymcmc = mcmc(mymodel, monitors, moves)
mymcmc.burnin(1E5,100)
mymcmc.run(1E6)
```

**Listing 5:** Excerpt from the RevBayes script for *Step 1* of the stepwise Bayesian inference on the Bayesian relaxed clock example.

##### S2.2.2 Step 2

*Step 2* of the Bayesian stepwise inference takes the samples from the importance distribution generated in *step 1* as data. We achieve this using the function `readTreeTrace`. How many samples are actually taken is controlled by the thinning argument, that is, when we performed our tests using a different number of samples $N$ from *step 1* we simply applied a different thinning of the samples. The parameters and distributions for *step 2* are shown in Table S6 (see also Figure 3c). Compared to the joint inference, we do not have the layer including the data. Note that we use `dnEmpiricalSample` to represent the probability distribution for the importance sample. This `dnEmpiricalSample` is completely generic and can be used for any probability distribution.

**Table S6: Model parameter names and prior distributions for *step 2* of the stepwise Bayesian inference for the Bayesian relaxed clock example.**

| Parameter | $X$ | $f(X)$ |
|---|---|---|
| Mean clock rate | $m_c$ | Uniform$(0, 10^4)$ |
| Standard deviation of clock rates | $m_{sd}$ | Uniform$(0, 10^4)$ |
| Phylogeny | $\Psi$ | UniformTimeTree$(root)$ |

```
treetrace = readTreeTrace("output/step1.trees",
                          treetype="non-clock",
                          burnin=0.001,
                          thin=1000)

clock_rate_mean ~ dnUniform(0,1E4)
clock_rate_sd ~ dnUniform(0,1E4)

clock_rate_ln_mean := ln(clock_rate_mean)

branch_rate_prior = dnLognormal(clock_rate_ln_mean, clock_rate_sd)

time_tree ~ dnUniformTimeTree(rootAge=ROOT_AGE, taxa=taxa)
time_tree.setValue( true_tree )
```

```

root_branch_fraction ~ dnBeta(1,1)

phis ~ dnEmpiricalSample(
                  dnBranchRateTree( time_tree, branch_rate_prior, root_branch_fraction ) )
phis.clamp(treetrace.getTrees())


moves.append( mvScale( clock_rate_mean, weight=3 ) )
moves.append( mvScale( clock_rate_sd, weight=3 ) )
moves.append( mvNodeTimeSlideUniform(time_tree,weight=NUM_TAXA) )
moves.append( mvBetaProbability(root_branch_fraction, weight=2.0) )

monitors.append( mnFile(time_tree, filename="output/step2.trees", printgen=10) )

mymcmc.burnin(1E5,100)
mymcmc.run(1E6)
```

**Listing 6:** Excerpt from the RevBayes script for *Step 2* of the stepwise Bayesian inference on the Bayesian relaxed clock example.

#### S2.3 Results

For completeness, we present here the results from the toy example showing (a) more observations, and (b) posterior distributions for the mean parameter $\mu$.

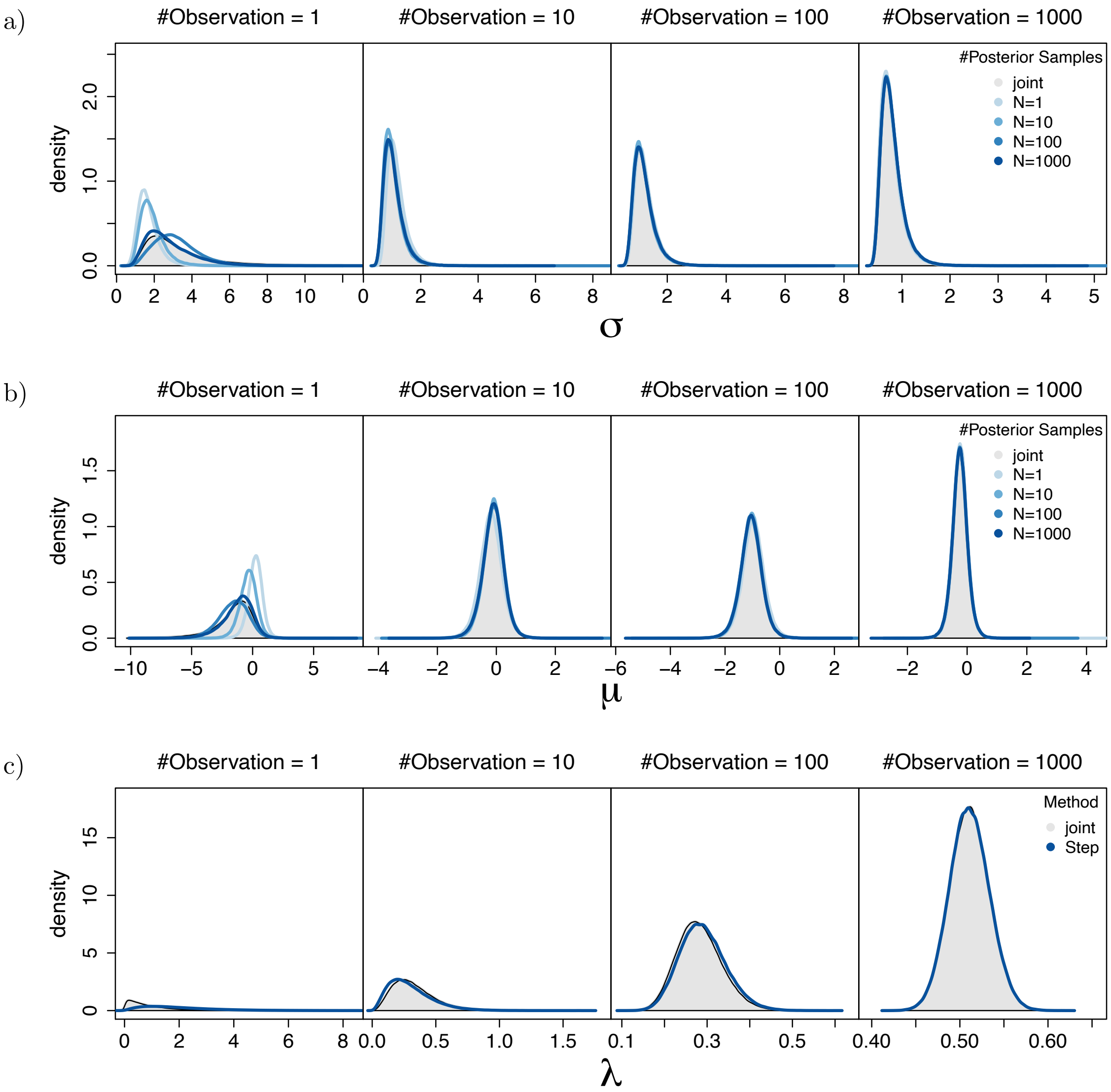


**Figure S1:** Estimated posterior distributions for the simple toy example as shown in Figure 1. In each row, we show the posterior distributions for different numbers of observations (columns) and different number of samples used in *step 2* (colors). The top row shows the posterior distributions of $\sigma$ and the middle row the posterior distribution of $\mu$. The bottom row shows the posterior and importance distribution of an arbitrarily picked $\lambda_i$. We observe that with few samples the joint and stepwise inferences disagree (left column; top row) which is due to divergence of the posterior distribution and importance distribution (left column; bottom row). Joint and stepwise inference are identical for many observations (right column). More samples $M$ of the importance distribution in *step 2* are beneficial but have a smaller impact than the number of observations.

### References

[1] TH Jukes and CR Cantor. Evolution of protein molecules. *Mammalian Protein Metabolism*, 3: 21–132, 1969.

[2] A. J. Drummond, S. Y. W. Ho, M. J. Phillips, and A. Rambaut. Relaxed Phylogenetics and Dating with Confidence. *PLoS Biology*, 4(5):e88, 2006.
